## Supplementary Figures and Tables for "Interplay between T3SS effectors, ExoY activation, and cGMP signaling in *Pseudomonas aeruginosa* infection"

A

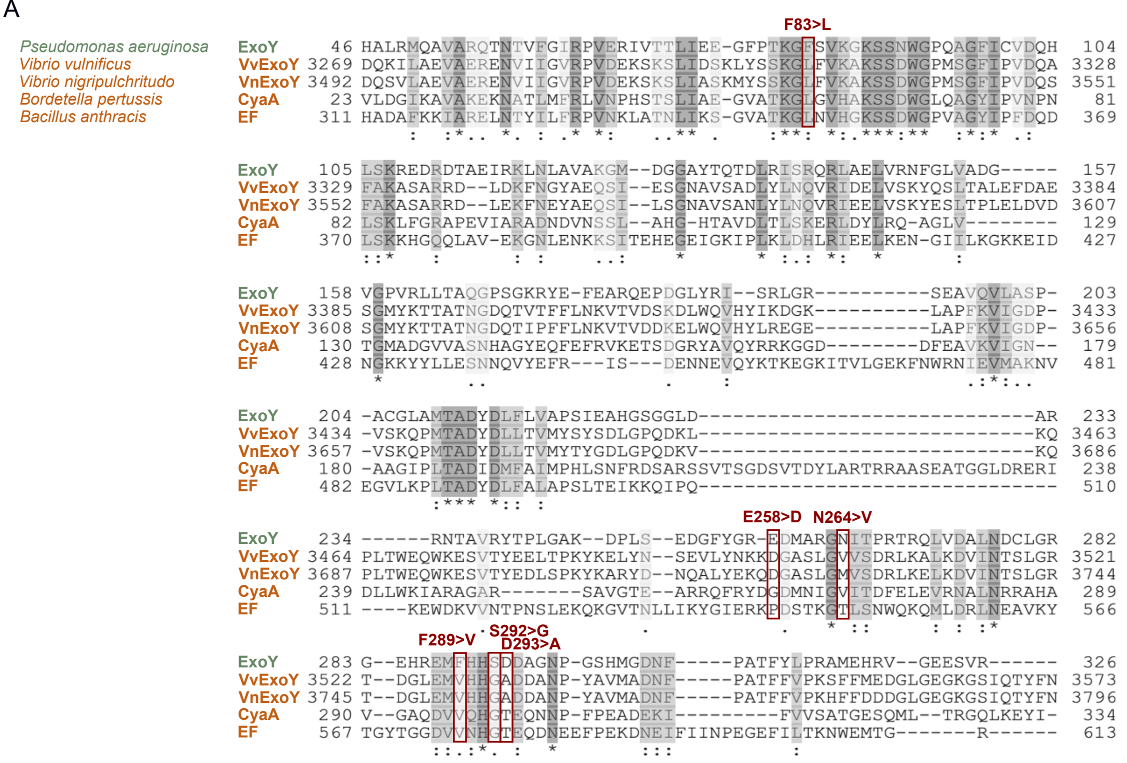

**Figure S1: Alignment of cAMP-inducing toxins with ExoY effector from *Pseudomonas aeruginosa*.**

**A.** Multiple sequence alignment of adenylate cyclase toxins from *Bordetella pertussis* and *Bacillus anthracis* (CyaA and EF respectively) and the ExoY-like modules from *Vibrio vulnificus* and *V. nigripulchritudo* (VvExoY and VnExoY respectively) with ExoY effector from *Pseudomonas aeruginosa* using the Clustal. Residues have been shaded to indicate different levels of conservation, with strictly conserved amino acids shown in dark gray and similar amino acids shown in lighter gray. A residue missing in one effector relative to the others is represented by "-". An asterisk "*" indicates that the residue is fully conserved among the effectors. Two dots ":" indicate a conserved substitution while one dot "." indicates semi-conserved substitutions. Numbers at the beginning and end of each sequence denote the amino acid positions. The four mutations selected in the nucleotide binding pocket of ExoY to change its substrate specificity are indicated in red.

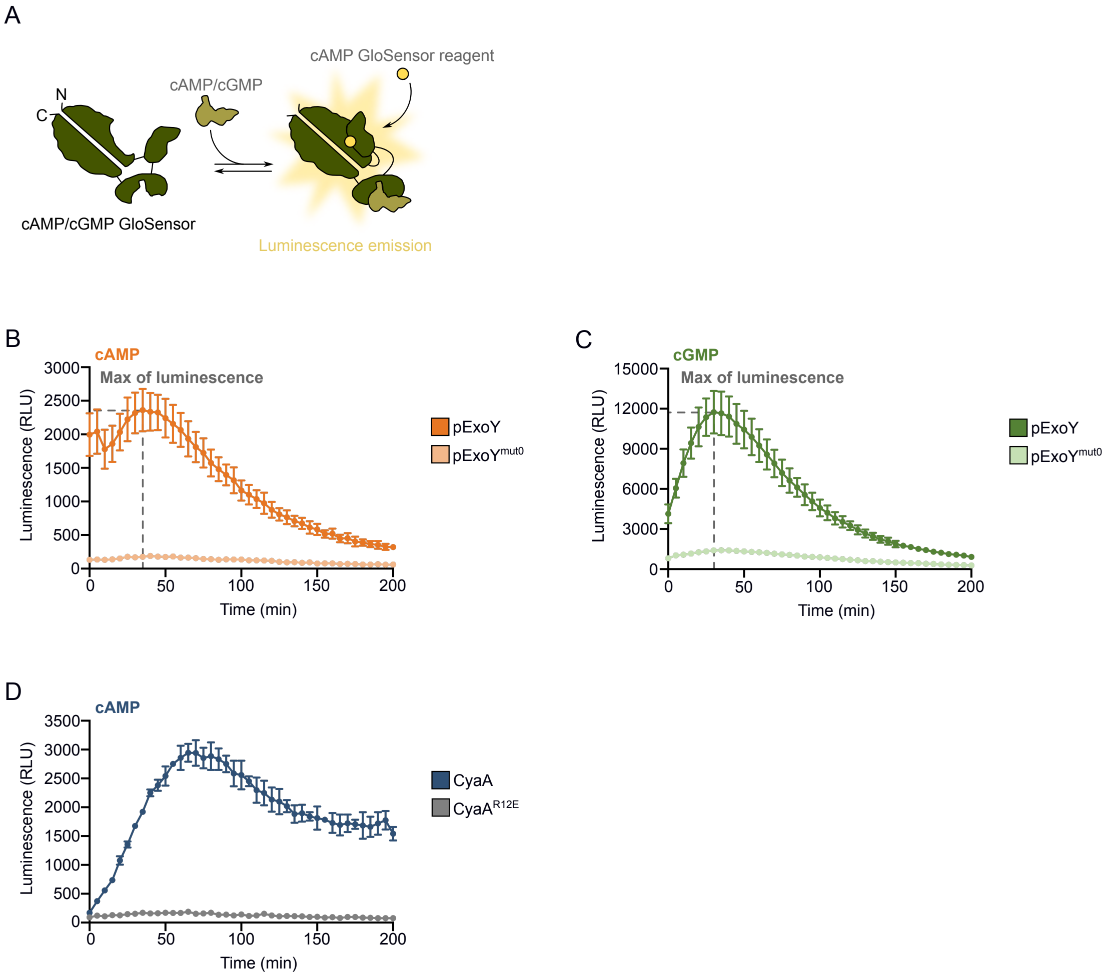

**Figure S2: Luminescence emission from cAMP- and cGMP-reporter cell lines.**

**A.** Representation of cAMP or cGMP Glosensor structure before and after interaction with second messengers. Binding of the analyte (cAMP or cGMP) induces conformational changes in the GloSensor, which emits luminescence following addition and binding of the Glosensor cAMP reagent. The magnitude of luminescence is proportional to analyte concentration. **B.** Increased luminescence emitted by stable clonal NCI-H292 cells expressing the cAMP- or **C.** cGMP Glosensor. Two days before acquisition, cells were co-transfected with plasmids encoding ExoY or catalytically inactive ExoY (ExoY$^{mut0}$) under control of the *TRE3G* promoter ($P_{TRE3G}$) and the transactivator required for activation. Prior to measurement, transfected cells were incubated with 1 µg/mL Doxycycline for 3h to trigger *exoY* expression. The cell medium was then replaced with fresh cell medium supplemented with 5% Glosensor cAMP reagent. Acquisitions were performed at 24°C every 5 minutes for 200 minutes with an integration time of 1000 ms. The dashed line indicates the maximum luminescence values as reported in Figure 1 F and G. **D.** Increased luminescence emitted by stable clonal cell line expressing the cAMP Glosensor after intoxication with 1 nM of CyaA WT or CyaA$^{R12E}$ proteins. Recombinant CyaA$^{R12E}$ protein does not translocate and was used as a negative control[2]. Cells were incubated for 1 h in CyaA activation medium supplemented with 5% Glosensor cAMP reagent prior to the addition of CyaA. Acquisitions were performed at 24°C every 3 minutes for 200 minutes with an integration time of 1000 ms. Error bars indicate SD.

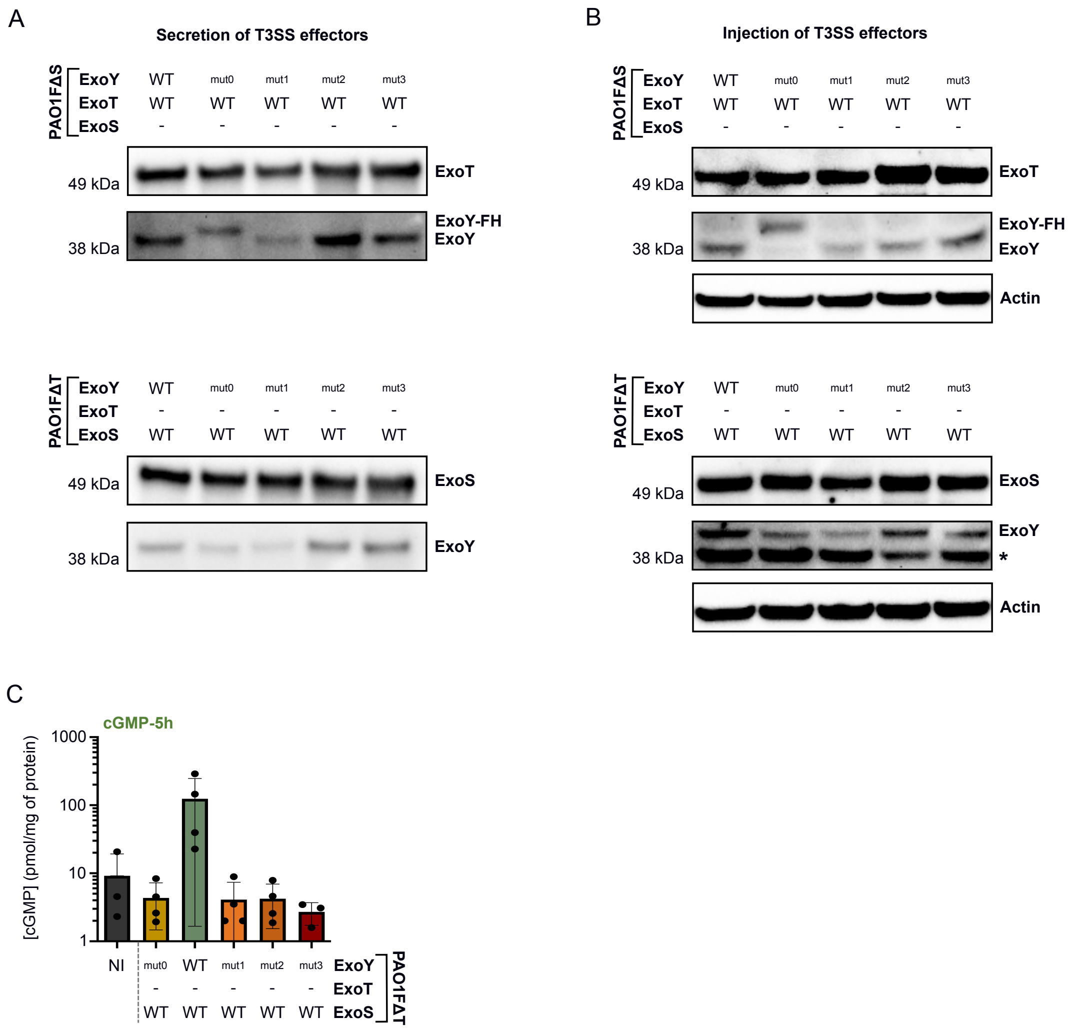

**Figure S3: Analysis of secretion and injection of T3SS effectors and cGMP production by PAO1FΔS or PAO1FΔT strains.**

**A.** Western blot (WB) of secreted T3SS effectors in bacterial culture supernatants. Overnight cultures of PAO1FΔS and PAO1FΔT strains were diluted to 0.05 and grown for 3 h in LB supplemented with 5 mM EGTA and 20 mM $MgCl_2$ to secrete ExoT/ExoS and the different ExoY variants. The ExoY$^{mut0}$ variant in the PAO1FΔS strain has a higher molecular weight because it contains a Flag-His tag at its C-terminus. **B.** WB showing T3SS effectors injected into NCI-H292 cell line after infection for 5h at an MOI of 20 with PAO1FΔS and PAO1FΔT strains. The ExoY$^{mut0}$ variant in the PAO1FΔS strain has a higher molecular weight because it contains a Flag-His tag at its C-terminus. T3SS effectors were revealed with anti ExoS, ExoT or ExoY antibodies. β-actin was used as loading control. (*) represents a non-specific band corresponding to a cellular protein that reacts with our ExoY antibody, as this band is also visible in the uninfected condition. **C.** Quantification of cGMP production in NCI-H292 cells by ELISA after 5h of infection at MOI 20 with PAO1FΔT strains co-injecting ExoS with an ExoY variant.

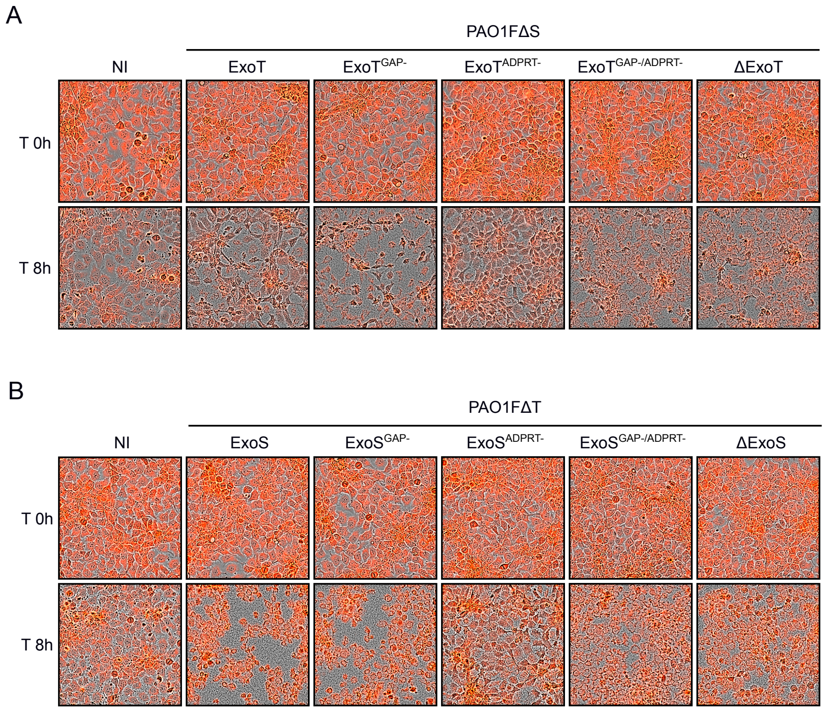

**Figure S4: Visualization of cell retraction induced by ExoT, ExoS or their derivatives.**
**A.** Images showing cell rounding induced by ExoT activities. Cells were incubated with 1 µM Cytotrace to label the cell cytoplasm and infected at MOI 50 for 8h with PAO1FΔS strains injecting ExoT toxin or its derivatives. Representative images show merged acquisition of phase contrast and Cytotrace fluorescence (red) at the beginning (T 0h) and after 8 hours of infection (T 8h). **B.** Representative images of cell rounding induced by ExoS activities. Similar to **A.** but cells were infected at MOI 20 for 8h with PAO1FΔT strains injected ExoS toxin or its derivatives.

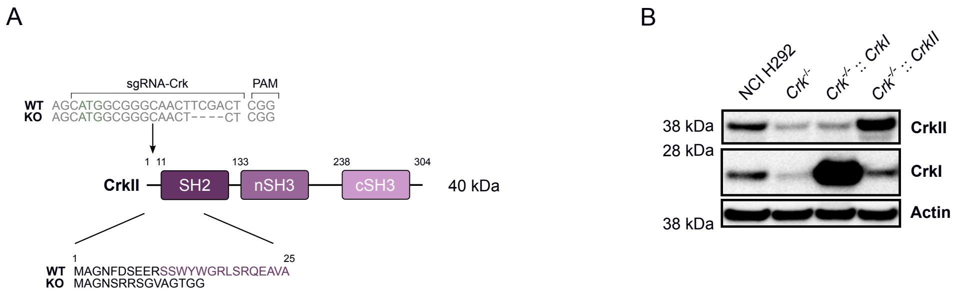

**Figure S5: Analysis of phospho-CrkII and Crk family proteins expression in native and Crk-deficient NCI-H292 cells.**

**A.** Alignments of native *crkII* sequence (WT) and mutant allele (KO) at the level of DNA and protein sequence showing four missing nucleotides in the mutant allele. The start codon is represented in green. The PAM sequence and the sequence of the sgRNA used for inactivation are shown. The mutant allele results in the expression of a truncated protein due to the introduced frame shift. The result is expected to be the same for CrkI as the deletion is at the beginning of the *crk* gene. **B.** WB of NCI-H292 and monoclonal Crk-/- NCI-H292 complemented or not with plasmids expressing CrkI or CrkII. The Crk proteins were revealed with the mouse recombinant monoclonal Crk antibody, targeting both proteins. β-actin was used as loading control.

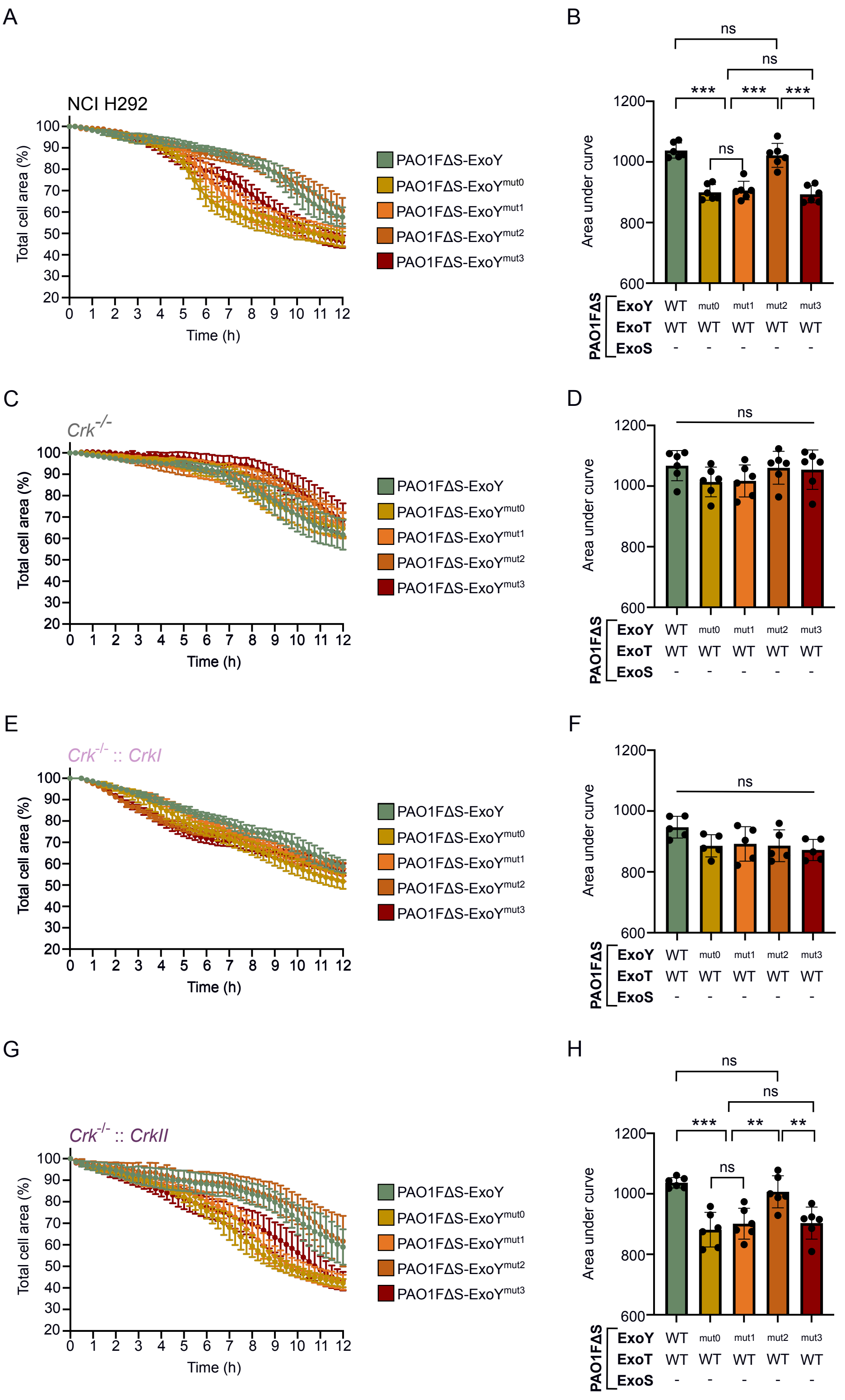

**Figure S6: ExoT-induced cell retraction in NCI-H292 WT or Crk-deficient cells, complemented or not with CrkI or CrkII protein.**

**A.** ExoT-induced cell retraction over time in NCI-H292 cells, **C.** monoclonal Crk-deficient cells, **E.** monoclonal Crk-deficient cells complemented with CrkI, and **G.** monoclonal Crk-deficient cells complemented with CrkII. Cytotrace was used to label the cell cytoplasm and cells were infected at MOI of 20 with PAO1FΔS strains co-injecting ExoT with an ExoY variant. Intoxication was recorded every 15 min by time-lapse microscopy. Results are represented as the mean percentage (±SD) of total cell area. Panels **B.**, **D.**, **F.**, and **H.** correspond to quantifications of the area under the curve (AUC) for each infection curve. The lower the AUC, the more cytotoxic the strain. Each dot represents a separate cell retraction assay. Statistical differences were established by one-way ANOVA ($p < 0.001$), followed by Tukey's test (*$p = 0.033$, **$p = 0.002$, ***$p < 0.001$, n.s., non-significant)

Supplementary Table 1 : Strains and plasmids

| Names | Description | Reference |
|---|---|---|
| ***E. coli* strains** | | |
| DH5α | F- Φ80*lac*ZΔM15 Δ(*lac*ZYA-argF) U169 *rec*A1 *end*A1 *hsd*R17($r_k$-, $m_k$+) *pho*A *sup*E44 *thi*-1 *gyr*A96 *rel*A1 λ- | Invitrogen |
| BLR | *E. coli* BL21, *recA* $^-$ : F$^-$*ompThsdS$_B$*($r_B$ $^-$$m_B$$^-$ ) *gal dcm* Δ(srl-recA)306::Tn10 (Tet$^R$ ) | Novagen |
| S17.1 (λ pir$^+$) | *pro*, *res*$^-$ *hsd*R17 ($r_k$-, $m_k$+) *rec*A$^-$ with an integrated RP4-2-Tc::Mu-Km::Tn7, λ pir$^+$, Tp$^r$ | [1] |
| ***P. aeruginosa* strains** | | |
| PAO1F (RP1831) | Secretion of ExoS, ExoT and ExoY toxins | [2] |
| PAO1FΔS (RP1883) | Secretion of ExoT and ExoY toxins | [3] |
| PAO1FΔS-ExoY$^{mut0}$ | As PAO1FΔS, but secretion of ExoT toxin with a catalytically inactive ExoY toxin (ExoY$^{K81M-K88I}$-FH) | This study |
| PAO1FΔS-ExoY$^{mut1}$ | As PAO1FΔS, but secretion of ExoT toxin with ExoY toxin carrying N264V, F289V, S292G and D293A mutations (ExoY$^{mut1}$) | This study |
| PAO1FΔS-ExoY$^{mut2}$ | As PAO1FΔS, but secretion of ExoT toxin with ExoY toxin carrying F83L and S292G mutations (ExoY$^{mut2}$) | This study |
| PAO1FΔS-ExoY$^{mut3}$ | As PAO1FΔS, but secretion of ExoT toxin with ExoY toxin carrying F83L, E258D and S292G mutations (ExoY$^{mut3}$) | This study |
| PAO1FΔS-ExoT$^{GAP-}$ | As PAO1FΔS, but secretion of ExoY toxin with ExoT toxin carrying R149K mutation to inactivate its Rho-GAP activity (ExoT$^{GAP-}$) | This study |
| PAO1FΔS-ExoT$^{ADPRT-}$ | As PAO1FΔS, but secretion of ExoY toxin with ExoT toxin carrying E383D and E385D mutations to inactivate its ADPRT activity (ExoT$^{ADPRT-}$) | This study |
| PAO1FΔS-ExoT$^{GAP-/ADPRT-}$ | As PAO1FΔS-ExoT$^{GAP-}$, but with ExoT$^{GAP-}$ toxin carrying E383D and E385D mutations in addition to inactivate its ADPRT activity (ExoT$^{GAP-/ADPRT-}$) | This study |
| PAO1FΔS-ExoT$^{ADPRT_ExoS}$ | As PAO1FΔS, but secretion of ExoY toxin with ExoT toxin whose ADPRT domain is replaced by that of ExoS | This study |
| PAO1FΔS-ExoY$^{mut0}$-ExoT$^{ADPRT_ExoS}$ | As PAO1FΔS-ExoY$^{mut0}$, but with ExoT toxin whose ADPRT domain is replaced by that of ExoS | This study |
| PAO1FΔS-ExoY$^{mut1}$-ExoT$^{ADPRT_ExoS}$ | As PAO1FΔS-ExoY$^{mut1}$, but with ExoT toxin whose ADPRT domain is replaced by that of ExoS | This study |
| PAO1FΔS-ExoT$^{GAP-_ADPRT_ExoS}$ | As PAO1FΔS-ExoT$^{GAP-}$, with ExoT$^{GAP-}$ toxin whose ADPRT domain is replaced by that of ExoS | This study |
| PAO1FΔT (RP1945) | Secretion of ExoS and ExoY toxins | [3] |
| PAO1FΔT-ExoY$^{mut0}$ | As PAO1FΔT, but secretion of ExoS toxin with a catalytically inactive ExoY toxin (ExoY$^{K81M-K88I}$) | This study |

| | | |
|---|---|---|
| PAO1FΔT-ExoY$^{mut1}$ | As PAO1FΔT, but secretion of ExoS toxin with ExoY toxin carrying N264V, F289V, S292G and D293A mutations (ExoY$^{mut1}$) | This study |
| PAO1FΔT-ExoY$^{mut2}$ | As PAO1FΔT, but secretion of ExoS toxin with ExoY toxin carrying F83L and S292G mutations (ExoY$^{mut2}$) | This study |
| PAO1FΔT-ExoY$^{mut3}$ | As PAO1FΔT, but secretion of ExoS toxin with ExoY toxin carrying F83L, E258D and S292G mutations (ExoY$^{mut3}$) | This study |
| PAO1FΔT-ExoS$^{GAP-}$ | As PAO1FΔT, but secretion of ExoY toxin with ExoS toxin carrying R146K mutation to inactivate its Rho-GAP activity (ExoS$^{GAP-}$) | This study |
| PAO1FΔT-ExoS$^{ADPRT-}$ | As PAO1FΔT, but secretion of ExoY toxin with ExoS toxin carrying E379D and E381D mutations to inactivate its ADPRT activity (ExoS$^{ADPRT-}$) | This study |
| PAO1FΔT-ExoS$^{GAP-/ADPRT-}$ | As PAO1FΔT-ExoS$^{GAP-}$, but with ExoS$^{GAP-}$ toxin carrying E379D and E381D mutations in addition to inactivate its ADPRT activity (ExoS$^{GAP-/ADPRT-}$) | This study |
| PAO1FΔST (RP1947) | Secretion of ExoY effector | [3] |
| PAO1FΔSTY (RP1949) | Secretion of none of the T3SS effectors | [4] |
| **Bacterial plasmids** | | |
| pUM460 | Plasmid for temperature-inducible expression of ExoY protein with a C-terminal Flag-His tag (ExoY-FH), [AmpR] | [5] |
| p1682 | Myc-PaExoY$^{K81M-K88I}$ in YEpGal555, [AmpR] | [6] |
| pB26 | As pUM460, but ExoY$^{K81M-K88I}$-FH (ExoY$^{mut0}$-FH) | This study |
| pUM717 | As pUM460, but ExoY$^{N264V-F289V-S292G-D293A}$-FH (ExoY$^{mut1}$-FH) | This study |
| pUM721 | As pUM460, but ExoY$^{F83L}$ -FH | This study |
| pUM727 | As pUM460, but ExoY$^{F83L-S292G}$ -FH (ExoY$^{mut2}$-FH) | This study |
| pUM733 | As pUM460, but ExoY$^{F83L-E258D-S292G}$-FH (ExoY$^{mut3}$-FH) | This study |
| pEX18Gm | Allelic exchange vector to introduce mutations in *P. aeruginosa* strains, containing *sacB* gene, a conditional lethal gene conferring sucrose sensitivity, [GmR] | [7] |
| pB93Gm | As pEX18Gm but with *exoY* gene from PAO1F strain and its surrounding sequences, fused to FH and containing the PstI restriction site | This study |
| pUM549 | As pB93Gm, but with the K81M and K88I mutations (mut0) in the *exoY* gene | This study |
| pUM707 | As pEX18Gm, but with *exoY* gene from PAO1F strain and its surrounding sequences | This study |
| pUM712 | As pUM707 but containing the N264V, F289V, S292G and D293A mutations (mut1) in the *exoY* gene | This study |
| pUM728 | As pUM707 but containing the F83L and S292G mutations (mut2) in the *exoY* gene | This study |
| pUM736 | As pUM707 but containing the F83L, E258D and S292G mutations (mut3) in the *exoY* gene | This study |

| | | |
|---|---|---|
| pUM737 | As pEX18Gm but with the GAP domain of the PAO1F *exoS* gene and the R146K mutation ($GAP^-$) | This study |
| pUM738 | As pEX18Gm but with the ADPRT domain of the PAO1F *exoS* gene and the E379D and E381D mutations ($ADPRT^-$) | This study |
| pUM739 | As pEX18Gm but with the GAP domain of the PAO1F *exoT* gene and the R149K mutation ($GAP^-$) | This study |
| pUM740 | As pEX18Gm but with the ADPRT domain of the PAO1F *exoT* gene and the E383D and E385D mutations ($ADPRT^-$) | This study |
| pUM752 | pEX18Gm-derived vector used to replace the ADPRT domain of ExoT with that of ExoS in the *exoT* gene of PAO1F strains | This study |
| **Mammalian plasmids** | | |
| pAcGFP-N1 | Expression vector for fusing AcGFP1 to the C-terminus of a partner protein under control of $P_{CMV\ IE}$ | Clontech |
| pTRE3G-BI | Tet-On 3G bidirectional inducible expression plasmid | TakaraBio |
| pVR2 | As pTRE3G-BI, but with AcGFP1 on MCS1 | This study |
| pUM445 | pBAD18 with ExoY-His | [5] |
| pUM542 | As pTRE3G-BI but with AcGFP1 on MCS1 and ExoY on MCS2 | This study |
| pUM546 | As pTRE3G-BI but with AcGFP1 on MCS1 and $ExoY^{K81M-K88I}$ ($ExoY^{mut0}$) on MCS2 | This study |
| pUM700 | As pTRE3G-BI but with AcGFP1 on MCS1 and $ExoY^{N264V-F289V-S292G-D293A}$ ($ExoY^{mut1}$) on MCS2 | This study |
| pUM731 | As pTRE3G-BI but with AcGFP1 on MCS1 and $ExoY^{F83L-S292G}$ ($ExoY^{mut2}$) on MCS2 | This study |
| pUM734 | As pTRE3G-BI but with AcGFP1 on MCS1 and $ExoY^{F83L-E258D-S292G}$ ($ExoY^{mut3}$) on MCS2 | This study |
| pGloSensor™-22F cAMP | Transfection plasmid containing a biosensor measuring cAMP levels in cells by luminescence | Promega |
| pGloSensor™-40F cGMP | Transfection plasmid containing a biosensor measuring cGMP levels in cells by luminescence | Promega |
| pLVX-IRES-neo | HIV-1-based lentiviral expression vector, [AmpR/NeoR/KanR] | TakaraBio |
| pLVX-neo-cAMP | As pLVX-IRES-neo, but with cAMP GloSensor | This study |
| pLVX-neo-cGMP | As pLVX-IRES-neo, but with cGMP GloSensor | This study |
| psPAX2 | Lentiviral packaging plasmid | Addgene |
| pMD2.G | VSV-G envelope expressing plasmid | Addgene |
| pUM517 | As pAcGFP1-N1, but ExoY-AcGFP1 | This study |
| pUM518 | As pAcGFP1-N1, but $ExoY^{K81M}$-AcGFP1 | [5] |
| pEF1α | Doxycycline-inducible mammalian expression vector allowing Tet-On 3G transactivator expression, [AmpR/NeoR/KanR] | TakaraBio |

| | | |
|---|---|---|
| pLentiCRISPRv2 | Lentiviral vector containing Cas9 nuclease gene, [AmpR/PuroR] | Addgene |
| pTwist Amp_CrkI | Twist cloning vector with a pMB1 origin and containing the modified *CrkI* gene, [AmpR] | This study |
| pTwist Amp_CrkII | Twist cloning vector with a pMB1 origin and containing the modified *CrkII* gene, [AmpR] | This study |
| pLVX-neo_CrkI | As pLVX-IRES-neo, but with the modified *CrkI* gene | This study |
| pLVX-neo_CrkII | As pLVX-IRES-neo, but with the modified *CrkII* gene | This study |

Supplementary Table 2 : List of primers

| Names | Sequence (5' → 3’) |
|---|---|
| **Primers for mutations*** | |
| VD5-F83L_F | GGTTTCCCGACCAAGGGCCTGTCGGTGAAGGGGAAAAG |
| VD5-F83L_R | CTTTTCCCCTTCACCGACAGGCCCTTGGTCGGGAAACC |
| VD11-S292G_F | GGAGATGTTTCACCACGGCGACGATGCGGGCAAC |
| VD11-S292G_R | GTTGCCCGCATCGTCGCCGTGGTGAAACATCTCC |
| VD13-E258D_F | CGGATTCTATGGCAGGGATGATATGGCCAGGGGAAAC |
| VD13-E258D_R | GTTTCCCCTGGCCATATCATCCCTGCCATAGAATCCG |
| VD-N264V_F | GGCCAGGGGAGTCATCACTCCGC |
| VD-N264V_R | GCGGAGTGATGACTCCCCTGGCC |
| VD-F289V-S292G-D293A_F | CAGGGAGATGGTTCACCACGGCGCCGATGCGGGCA |
| VD-F289V-S292G-D293A_R | TGCCCGCATCGGCGCCGTGGTGAACCATCTCCCTG |
| ExoSR146K-Up_F | CAACAAGCTTCATACCTTGGTCGATCAGCTTTTGTC (HindIII) |
| ExoSR146K-Up_R | GGAGATGGGGCGCTGAAATCGCTGAGCACCGCCTT |
| ExoSR146K-Down_F | AGGCGGTGCTCAGCGATTTCAGCGCCCCATCTCC |
| ExoSR146K-Down_R | AAAAGAATTCCATCAGGAGAAGGCAACCATCATGC (EcoRI) |
| ExoSE379/381D-Up_F | CAACAAGCTTGACAGAAGCAGAGCCGTTTCGTC (HindIII) |
| ExoSE379/381D-Up_R | TCGAACTACAAGAATGATAAAGATATTCTCTATAACAAAGAGACCG |
| ExoSE379/381D-Down_F | CGGTCTCTTTGTTATAGAGAATATCTTTATCATTCTTGTAGTTCGA |
| ExoSE379/381D-Down_R | CAAAGAATTCGCGGATGCGGAAAAGTACCTG (EcoRI) |
| ExoTR149K-Up_F | GGGGAAGCTTCAGGAGACGTCAATCATCATGCATATTC (HindIII) |
| ExoTR149K-Up_R | GCGGTGGCCAGCGATTTCAGAGCGCCGTCGC |
| ExoTR149K-Down_F | GCGACGGCGCTCTGAAATCGCTGGCCACCGC |
| ExoTR149K-Down_R | CAACGAATTCCATGCCCCGGTCGATCAACGCC (EcoRI) |
| ExoTE383/385D-Up_F | GAAGAAGCTTGAGAGCGAGGTAAAGGGCGAGCC (HindIII) |
| ExoTE383/385D-Up_R | CTTGTCGTAGAGGATATCCTGATCATCGCCCTCGATC |
| ExoTE383/385D-Down_F | GATCGAGGGCGATGATCAGGATATCCTCTACGACAAG |
| ExoTE383/385D-Down_R | GAAGGGATCCCATCAATAGGCCATCTCGGTGGG (BamHI) |
| **Primers for PCR** | |
| PAO1-Up_F | GGGGAAGCTTGTTCACCGCTCATCACCTCGC |
| PAO1-Down_R | CGGCGAATTCGTTCTACCAGGAGATCGGCC |
| VD17_F | TCGGCCGACAAGGCGCTGG |
| VD17_R | GTCCGTCTCTGCCGTCGACCGGTCAGGCCAGATCAAGGCCG |
| VD18_F | GGAAGGATCCATGCATATTCAATCATCTCAGCAG (BamHI) |
| VD18_R | CCAGCGCCTTGTCGGCCGACTCGCCCTTTACCTCGCTC |
| VD19_F | CCGGTCGACGGCAGAGACGGAC |
| VD19_R | GGGAAAGCTTGGTGGGTGATGAGAATGGTC (HindIII) |
| Amplif-PAO1-ExoS_F | CCAGCCCGGAGAGACTGTTAATCG |
| Amplif-PAO1-ExoS_R | CAGGTGTTCTGGTTCACCACCCTG |
| Amplif-PAO1-ExoT_F | GTTCTCTTTCCGCGTGCTCCGAC |
| Amplif-PAO1-ExoT_R | CGAAATGGGGTGGCCACGATTTC |
| Amplif-PAO1-ExoY_F | CAGTTGCGCCATGTTCTTCG |

| | |
|---|---|
| Amplif-PAO1-ExoY_R | GACAAGGCATGAGCACCAGC |
| Amplif-PAO1-ExoS-ADPRT_F | GGAGCTGGATGCGGGACAAAAG |
| VD20_F | GAGCCATGGCGGGAAATTTTG |
| VD20_R | CCTCTCTTGCCTTCGCTGTC |
| PCR-Crk_F | CTGCCTAGAAGTCCCAGGTCG |
| PCR-Crk_R | CCAAACCCGGCTCTCCAACAC |
| UM345 | GAGAGCTAGCCACCATGGCGCGTATCGACGGTCATCGTCAG (NheI) |
| UM246 | GGGGCTCGAGGACCTTACGTTGGAAAAAGTCG (XhoI) |
| UM245 | GGGGAATTCACCATGCGTATCGACGGTCATCG (EcoRI) |
| UM374 | GGGGACGCGTTCAGACCTTACGTTGGAAAAAGTCGAGC (MluI) |
| #124 | TATAAAGCTTGTTCACCGCTCATCACCTCG (HindIII) |
| #125 | TATACTGCAGTTTCCCGCATATGCCAAAGAAG (PstI) |
| #126 | TATATCTAGAAGGTCGTTGGCAAAGCCCG (XbaI) |
| #127 | TATAGAATTCGTTCTACCAGGAGATCGGCCA (EcoRI) |
| #128 | TATACTGCAGCATGGGCCGTATCGACGG (PstI) |
| #129 | TATATCTAGAATCAGTGGTGGTGGTGGTGG (XbaI) |
| **Primers for sequencing** | |
| UM316 | GGCTCTACAGGACCTGTTC |
| M13uni | AGGGTTTTCCCAGTCACGACGTT |
| M13rev | GAGCGGATAACAATTTCACACAGG |
| pLVX-neo_Seq_F | GCCATCCACGCTGTTTTGACC |
| pLVX-neo_Seq_R | GCTTCGGCCAGTAACGTTAGG |
| Seq-PAO1-ExoY_F | CGCTGCCCGGATTCTTATTC |
| Seq-PAO1-ExoY_R | GCTTCGGAAAGTCCCGTTCG |
| Seq-PAO1-ExoS_F | CATTGCCCATGACCTTGAAGG |
| Seq-PAO1-ExoS_R | GTTTGCTTGCCAGGTCGAGAG |
| Seq-PAO1-ExoT_F | GCAAAAGGCACTGCCCCTG |
| Seq-PAO1-ExoT_R | TCCAGGGCATCGAGCAGTC |
| Seq3-ExoS_F | CTCTCGACCTGGCAAGCAAAC |
| Seq-Crk_F | CATGAATCCGCATGCGCAGTG |
| Seq-Crk_R | CTTTGTCCCCACGTTGCCTTG |
| **Oligonucleotides to knock-out *Crk* gene**$^{\Delta}$ | |
| sgRNA3-Crk_F | **CACCG**CATGGCGGGCAACTTCGACT |
| sgRNA3-Crk_R | **AAAC**AGTCGAAGTTGCCCGCCATG**C** |

* Red letters indicate mutations compared to the WT sequence

$^{\Delta}$ Bold letters represent overhangs for cloning in Esp3I-digested pLentiCRISPRv2

Supplementary Table 3: Modified *CrkI* and *CrkII* genes synthesized by Twist

| Names | Sequence (5'→ 3')* |
|---|---|
| Modified *CrkI* | TCTAGAGCCATGGCGGGAAATTTTGATTCTGAGGAGCGGAGTAGCTGGTACTG GGGGAGGTTGAGTCGGCAGGAGGCGGTGGCGCTGCTGCAGGGCCAGCGGCA CGGGGTGTTCCTGGTGCGGGACTCGAGCACCAGCCCCGGGGACTATGTGCTCA GCGTCTCAGAGAACTCGCGCGTCTCCCACTACATCATCAACAGCAGCGGCCC**TA** G**A**CC**T**CCGGTGCCACCGTCGCC**T**GCCCAGCCTCC**T**CC**T**GGGGTGAGCCCCTCCA GACTCCGAATAGGAGATCAAGAGTTTGATTCATTGCCTGCTTTACTGGAATTCTA CAAAATACACTATTTGGACACTACAACGTTGATAGAACCAGTTTCCAGATCCAG GCAGGGTAGTGGAGTGATTCTCAGGCAGGAGGAGGCGGAGTATGTGCGAGCC CTCTTTGACTTTAATGGGAATGATGAGGAAGATCTTCCCTTTAAGAAAGGAGAC ATCTTGAGAATCCGGGACAAGCCTGAAGAGCAGTGGTGGAATGCGGAGGACA GCGAAGGCAAGAGAGGGATGATTCCAGTCCCTTACGTCGAGAAGTATAGACCT GCCTCCGCCTCAGTATCGGCTCTGATTGGAGGTCGGTGAGGATCC |
| Modified *CrkII* | TCTAGAGCCATGGCGGGAAATTTTGATTCTGAGGAGCGGAGTAGCTGGTACTG GGGGAGGTTGAGTCGGCAGGAGGCGGTGGCGCTGCTGCAGGGCCAGCGGCA CGGGGTGTTCCTGGTGCGGGACTCGAGCACCAGCCCCGGGGACTATGTGCTCA GCGTCTCAGAGAACTCGCGCGTCTCCCACTACATCATCAACAGCAGCGGCCC**TA** G**A**CC**T**CCGGTGCCACCGTCGCC**T**GCCCAGCCTCC**T**CC**T**GGGGTGAGCCCCTCCA GACTCCGAATAGGAGATCAAGAGTTTGATTCATTGCCTGCTTTACTGGAATTCTA CAAAATACACTATTTGGACACTACAACGTTGATAGAACCAGTTTCCAGATCCAG GCAGGGTAGTGGAGTGATTCTCAGGCAGGAGGAGGCGGAGTATGTGCGAGCC CTCTTTGACTTTAATGGGAATGATGAGGAAGATCTTCCCTTTAAGAAAGGAGAC ATCTTGAGAATCCGGGACAAGCCTGAAGAGCAGTGGTGGAATGCGGAGGACA GCGAAGGCAAGAGAGGGATGATTCCAGTCCCTTACGTCGAGAAGTATAGACCT GCCTCCGCCTCAGTATCGGCTCTGATTGGAGGTAACCAGGAGGGTTCCCACCCA CAGCCACTGGGTGGGCCGGAGCCTGGGCCCTATGCCCAACCCAGCGTCAACAC TCCGCTCCCTAACCTCCAGAATGGGCCCATATATGCCAGGGTTATCCAGAAGCG AGTCCCCAATGCCTACGACAAGACAGCCTTGGCTTTGGAGGTCGGTGAGCTGG TAAAGGTTACGAAGATTAATGTGAGTGGTCAGTGGGAAGGGGAGTGTAATGG CAAACGAGGTCACTTCCCATTCACACATGTCCGTCTGCTGGATCAACAGAATCCC GATGAGGACTTCAGCTGAGGATCC |

* Red letters represent mutations that prevent the Cas9 endonuclease from cleaving the gene. Bold letters represent silent mutations aimed at reducing %GC content. The yellow and blue letters correspond to the XbaI and BamHI restriction sites respectively.
